# Green emitting carbon quantum dots (GCQDs) to probe endocytic pathways in cells; for tissue and *in vivo* bioimaging

**DOI:** 10.1101/2022.04.23.489248

**Authors:** Pankaj Yadav, Krupa Shah, Krupa Kansara, Subhajit Das, Ashutosh Kumar, Rakesh Rawal, Dhiraj Bhatia

## Abstract

Small sized, carbon-based organic nanoparticles have recently gained attention due their advantage of biocompatibility, photostability and biological non-toxicity as compared to their inorganic counterparts. Herein, a new class of small (5-8 nm), green emitting fluorescent carbon quantum dots (GCQDs) were synthesized using organic substrates like citric acid and ascorbic acid in aqueous solvent containing water and ethanol. The very small size and bright green photoluminescence prompted their use for both *in vitro* and *in vivo* bioimaging. GCQDs were uptaken via clathrin mediated pathways in mouse kidney and liver primary cells. Similarly, they showed active uptake and distribution in the zebrafish embryo model system. The optical tunability and surface modification properties of these GCQDs provide a platform to be explored for them to emerge as a new class of targeted bioimaging entities, as well as tools for biomedical applications.

## Introduction

Carbon based quantum dots (CQDs), discovered in 2004 in an experiment of purification of single walled carbon nanotubes revealed their excellent property of photoluminescence owing to their small size and quantum effects^1^. CQDs’ small size impart them exceptional electronic and optical properties, which makes them highly fluorescent^2,3,4,5,6^. Carbon based nanoparticles are biocompatible, non-toxic, photostable, environment friendly and have low cost of synthesis compared to heavy metal quantum dots^7^. The mentioned advantages make them highly desirable for biomedical applications such as targeted bioimaging, biosensing, drug delivery, gene delivery, and disease diagnosis^8^.

Green synthesis of CQDs has been applied using various carbon sources^9^ and these QDs have been explored for various biological applications in sensing, photocatalysis^10^, fluorescent patterning ink^11^, antibacterial^12^ and bioimaging^13^. Plants, pulp and fruit juices have been widely used for the synthesis of CQDs^9^. Citric acid and ascorbic acid are the main and primary components in the composition of these citrus fruits. Citric acid^14^ is being explored for synthesis of CQDs and tuning their fluorescence by doping with nitrogen and sulphur for LEDs^15^. Citric acid with DMF and ethylenediamine has been used to demonstrate QDs with applications in theranostics^16^ and printing ink ^17,18^. Common sources like lemon juice ^19^ and orange juice^20^ have been reported for synthesis of fluorescent CQDs respectively for bioimaging applications. Green fluorescent CQDs have been explored widely for sensing and bioimaging applications^21^ but their applications in biological systems are limited owing to the limited understanding of their interactions with cells and tissues. Knowing the biological behaviour i.e., binding to membranes^22^, cellular uptake, cellular fates, of CQDs is a crucial and critical step in designing nanoparticle-based therapeutics. The understanding of the internalization pathways of CQDs in cells and tissues offers a great advantage for development of therapeutic strategies for their future exploration. The knowledge of the endocytosis pathway has allowed the engineering of nanoparticles to deliver drugs for cancer therapy ^23,24,25,26^.

Keeping these aspects in focus, we asked in this study, if we can use the component of citrus fruit juices to synthesize CQDs with size tunability and emission in visible regions of the spectrum? If yes, can we use them to study their fluorescence and biological uptake in cells, tissues and whole organisms. Zebrafish is an attractive and ideal *in vivo* model for drug delivery and novel nano formulation uptake studies, due to their small size, transparency and rapid growth^27^. Previous study revealed that the size of nano formulations contributes to the major role for uptake analysis in zebrafish embryos^28^. The aim of the current study was to provide a holistic understanding of accumulation and uptake of green fluorescent CQDs in the ideal zebrafish embryo model.

Herein, we present a novel type of water soluble, green fluorescent CQDs (GCQDs) to investigate the endocytosis pathways in cells and uptake in tissues and *in vivo*. The GCQDs (5-8 nm) were synthesized using the main components of citrus fruit juice (citric acid and ascorbic acid). These GCQDs are non-toxic to cells and biocompatible in mammalian cells. These GCQDs are efficiently uptaken in different types of mammalian cells via clathrin mediated pathways. The endocytosis of these GCQDs was then successfully used for bioimaging endocytic pathways in mammalian cells such as mouse primary cells. Further, we show their uptake in tissues like kidneys and liver from mice as well as specific targeting in zebrafish embryos (**Figure 1**). Our new GCQDs present a novel class of bioimaging entities which could be further modified to fine-tune their structure and emission properties for targeted biomedical applications such as biosensing and therapeutics *in vivo*.

**Figure 1.**
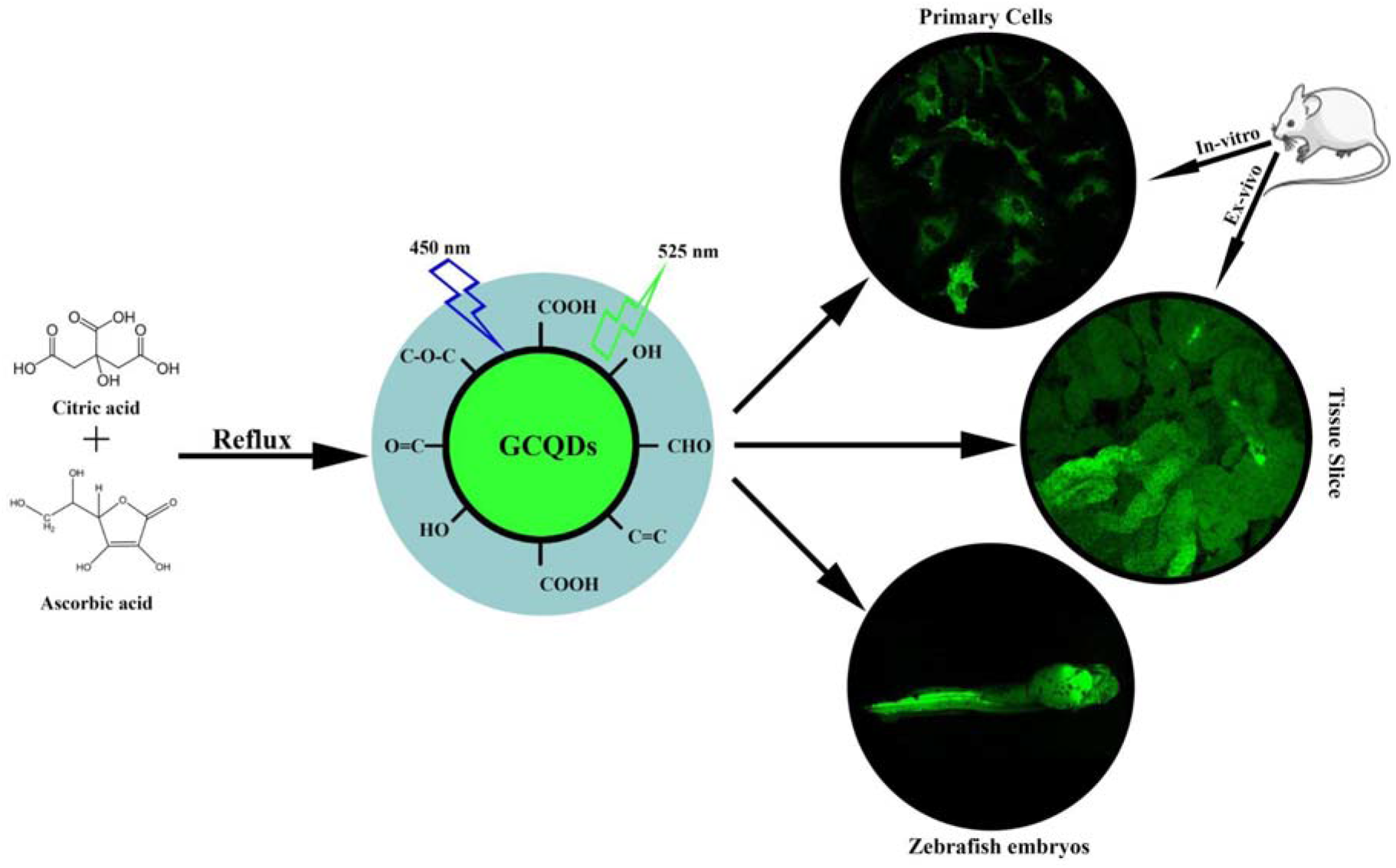
Synthesis and bioimaging applications of GCQDs. Citric and ascorbic acid are refluxed in ethanol-water mix to give bright green emitting carbon-based quantum dots, which can be used for bioimaging applications in cells, tissues and organisms like zebrafish embryos.

## Results and Discussions

The GCQDs were synthesized using citric acid (CA) and ascorbic acid (AA) in a solvent having water and ethanol in 2:1 ratio. The precursors were dissolved in the solvent and heated at 130ºC for 12 hours as shown in **figure S1**.The reaction was cooled down naturally to room temperature and GCQDs were collected by centrifugation and removing the solvent and lyophilizing the GCQDs.

The synthesized GCQDs were extensively characterized using multiple biophysical and spectroscopic methods. UV-visible absorbance spectroscopy of GCQDs gave a characteristic peak at 247 nm at pH 1.85, 267 nm at pH 7 and at 265 nm by the lyophilized GCQDs redissolved in water having pH 7 as shown in **figure 2a**. The band around 270 nm correspond to n-π* transition C=O bond. The photoluminescence spectra revealed that maximum emission intensity was observed at a wavelength of 525 nm when excited at 450 nm wavelength as shown in **figure 2b**. Fourier transform infrared spectroscopy (FT-IR) revealed the surface properties of the GCQDs as shown in **figure 2c**. The broad peak at 3232 cm^-1^ corresponds to -OH stretch and a shoulder peak at 2989 cm^-1^ correspond to sp^3^ -CH stretch. Similarly, the peak at 1722 cm^-1^ corresponds to carbonyl group(-C=O), 1566 cm^-1^ corresponds to C=C, 1390 cm^-1^ corresponds to sp^3^ -CH bend. While 1262 cm^-1^ and 1045 cm^-1^ corresponds to -C-O stretch and -C-O-C-stretch respectively. Thus, FTIR spectroscopy reveals the presence of hydroxyl and carboxyl groups which make the GCQDs highly water-soluble nanoparticles. GCQDs FTIR compared with citric acid and ascorbic acid shows that there is formation of new bonds revealed by new peaks at 3232cm^-1^, 2989 cm^-1^ and peaks shift at 1722 cm^-1^, 1566 cm^-1^, 1045 cm^-1^ in **figure 2d**. The powder XRD of GCQDs reveals three peaks at 11.32,21.89, 33.2 dictates the crystallinity of the GCQDs (**figure 2e**). The 1H NMR of GCQDs, peak between 1 ppm to 5 ppm shows the alcoholic hydrogen, and the peak between 1 ppm to 2 ppm is of sp^3^ H, between 2 ppm to 3 ppm is of carbonyl alpha H, between 3.3 ppm to 5 ppm is of alcohols, esters and ethers (**figure 2f**). The 13C NMR of GCQDs shows the carboxylic functional group having peaks from 160 ppm to 180 ppm, aldehyde with H and ketone with no H above 180 ppm. The peaks between 100 ppm to 120 ppm correspond to the C=C group, peaks between 40 ppm to 80 ppm show alcohols, esters and ethers with and without H. The peak between 10 ppm to 30 ppm shows sp^3^ carbon (**figure 2g**).The morphology (size and shape) of the GCQDs was revealed by TEM and AFM as shown in **figure 2h,i**. The morphology of the GCQDs seems to be spherical and appears crystalline in nature as shown in **figure 2h**. The size of the GCQDs was calculated to be 5-8 nm for 61% (19/31) of the counted nanoparticles and the similar size was confirmed using AFM as shown in **figure 2i**.

**Figure 2.**
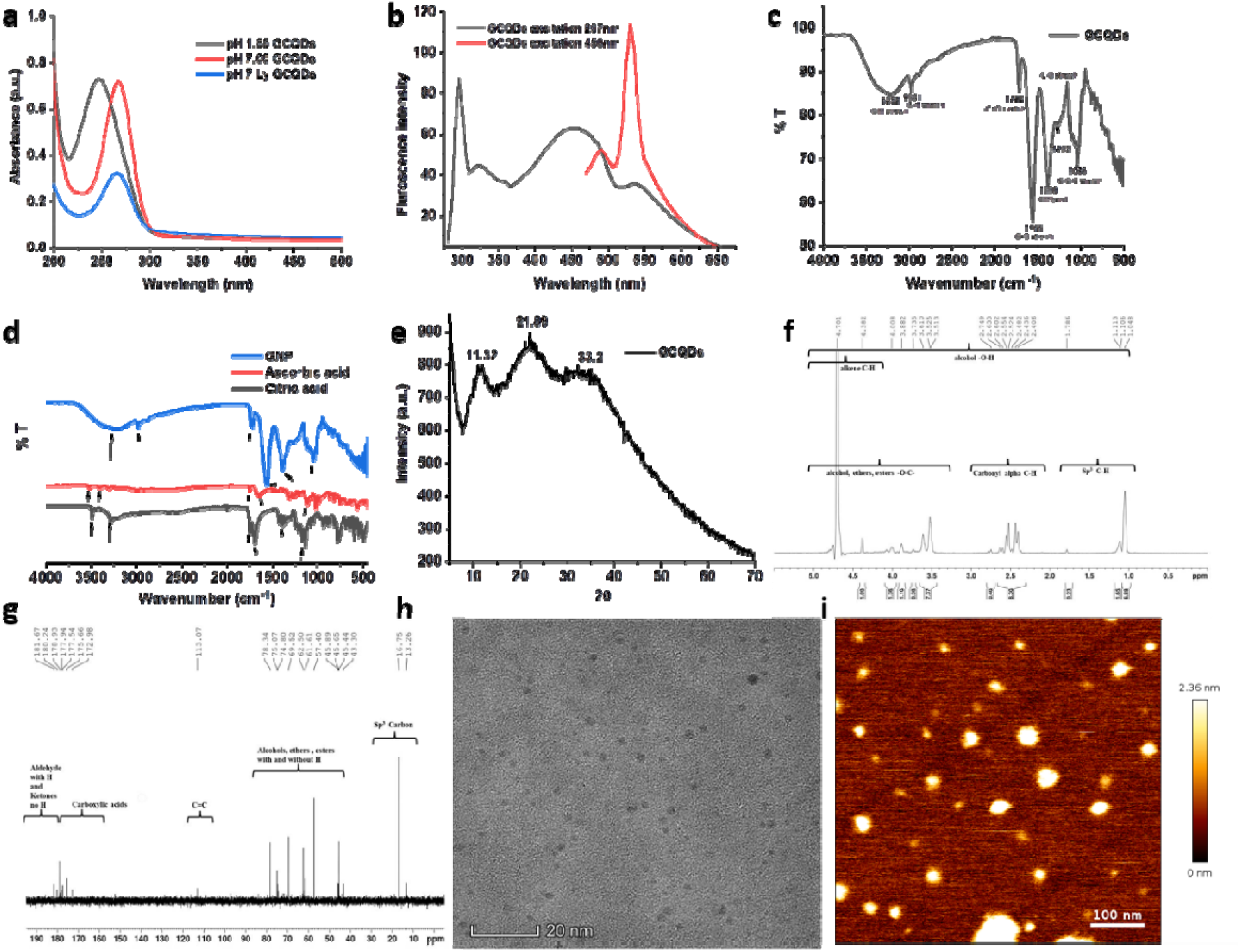
Characterization of GCQDs. **(a)** The UV visible spectra of GCQDs showed the absorbance at 247 nm for GCQDs at pH 1.85, at 268 nm for GCQDs at pH 7 and at 265 nm for lyophilized GCQDs dissolved in deionized water at pH 7 **(b)** The lyophilized GCQDs dissolved in deionized water at pH 7 shows several emission peaks 300nm, 325 nm, 450 nm, and 525 nm when excited at 250 nm, however GCQDs exhibited maximum emission at 525 nm when excited at a wavelength of 450 nm **(c)** FTIR Spectra of GCQDs shows a broad peak at 3232 cm^-1^ which corresponds to -OH stretch, the peak at 2981 cm^-1^ shows -CH stretch, the peak at 1722 cm^-1^ shows carbonyl group (-C=O), 1390 cm^-1^ shows -OH bend, 1262 cm^-1^ corresponds to -C-O stretch and 1045 cm^-1^ shows C-O-C stretch **(d)** FTIR spectra comparison of GCQDs with citric and ascorbic acid shows the shift in the peaks at 1744 cm^-1^ (in CA), 1992 cm^-1^ (of CA) to 1718 cm^-1^ and 1563 cm^-1^ (of GCQDs) respectively and while disappearance of bonds corresponding to peaks at 1143cm^-1^ (in CA), 775cm^-1^ (in CA) and 1111 cm^-1^ (in AS), 743 cm^-1^ (in AS), 3525 cm^-1^ (in AS) as compared to GCQDs. (**e**) The powder XRD of the lyophilized GCQDs at pH 7 shows peaks at 2 theta = 11.32, 21.89, 33.2 showing the crystalline nature of the GCQDs **(f)** NMR 1H spectra, the peaks between 1 ppm to 5 ppm shows the alcoholic hydrogen, and the peak between 1 ppm to 2 ppm is of sp^3^ H, between 2 ppm to 3 ppm is of carbonyl alpha H, between 3.3 ppm to 5 ppm is of alcohols, esters and ethers **(g)** NMR 13C spectra, peaks from 160 ppm to 180 ppm corresponds to carboxylic acid, peaks above 180 ppm shows aldehyde with H and ketone with no H. The peaks between 100 ppm to 120 ppm correspond to the C=C group, peaks between 40 ppm to 80 ppm show alcohols, esters and ethers with and without H. The peak between 10 ppm to 30 ppm shows sp^3^ carbon **(h)** TEM image of the GCQDs shoes size between 5-8 nm (count = 50) **(i)** AFM shows size distribution of the GCQDs, the bar graph shows that 60% of the GCQDs are in range of 5-8 nm (count 31).

### Cellular internalization and non-toxicity of GCQDs

To understand and study the applications of synthesized GCQDs in bio imaging as molecular probes and drug delivery, cellular cytotoxicity MTT assay was performed on epithelial RPE1 cells. MTT assay was performed with increasing concentration of GCQDs (50, 100, 150, 200, 250, 300, 350, 400 µg/ml) on RPE1 cells and cell viability was calculated after 12 hours of incubation. MTT assay revealed that, cell viability was negligibly affected at lower concentration 50 - 200 µg/ml, while percent cell viability decreased up to ∼60% as the concentration increased up to 350-400 µg/ml (**Figure 3a**), indicating that increasing the GCQDs concentration up to 300 µg/ml does not affect the overall cell viability which further suggesting the non-toxic effect of GCQDs to the overall cell survival and proliferation of RPE1 cells.

**Figure 3.**
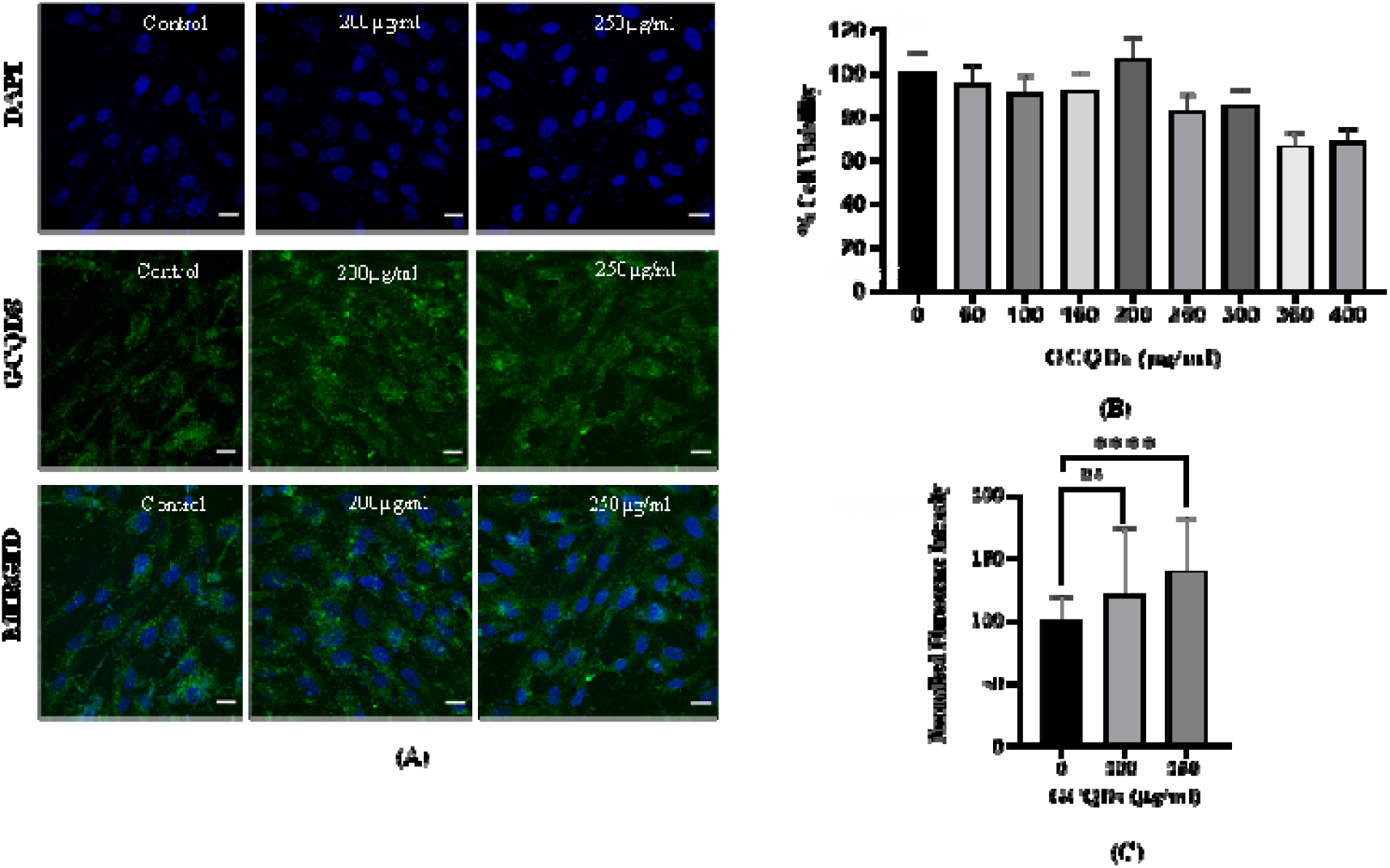
Uptake and cytotoxicity of GCQDs in epithelial RPE1 cells. **(A)** % cell viability of GCQDs after MTT assay. **(B)** Confocal Images of concentration dependent internalization of GCQDs in RPE1 cells. **(C)** Normalised fluorescence intensity and quantification analysis of different concentrations of GCQDs. Scale bar is set at 20 µm of all the images. **** denotes statistically significant p-value (p < 0.0001). whereas, ns indicates non-significant p-value. (One-way ordinary ANOVA test). Total 35 cells and n = 2 independent experiments were performed.

To understand the cellular uptake properties and internalization of developed GCQDs, a concentration dependent study was performed at specific time internal on RPE1 cells and analysed by confocal microscopy (**Figure 3b**,**c**). RPE1 cells were incubated with different concentrations of GCQDs (200, 250 and 300µg/ml) at 37ºC for 15 min in a 5% CO_2_ incubator and their cellular internalization, changes in cellular morphology and total fluorescence intensity were observed. Confocal analysis showed a successful internalization of GCQDs with a strong fluorescence response as the GCQDs concentration increased and morphology remained unchanged as the concentration increased.

### GCQDs endocytosed through clathrin mediated pathway in mouse kidney and liver derived primary cells

To substantiate the uptake and internalization of GCQDs for the particular cell type, we analysed the GCQDs uptake mode in mouse derived kidney and liver primary cells. Primary cell cultures derived from a small tissue piece and preparing a single cell suspension followed by enzymatic digestion represent a more realistic model than monoclonal cell lines, as their cell distribution and cell types are comparatively similar to solid tissue and represent physiologically more relevant biological systems than the differentiated cell lines.

Cells uptake various nutrients from extracellular space principally through two key endocytic pathways - clathrin dependent and independent. There are other pathways as well like micropinocytosis mediated by actin ruffles, etc. but are less efficient and involved in uptake of macromolecular structures like viruses, bacteria^29^. Using established markers and inhibitors for the key endocytic pathways, we explored the mechanisms of GCQDs internalization into the cells. Hence, these endocytosis pathways were further explored in kidney and liver primary cells to assess the cellular internalization of GCQDs into these cells. Cells were incubated with transferrin (a well-established marker for clathrin mediated endocytosis), galectin-3 (a protein that binds to glycosylated cargo on membrane and induce the biogenesis of clathrin independent endocytic vesicles), and GCQDs alone and in combination with pathway specific inhibitors pitstop-2 (CME inhibitor) and lactose (CIE inhibitor) for 15 min at 37ºC in a 5% CO_2_ incubator and analysed by confocal microscopy. Cells when treated with CME pathway blocking inhibitor pitstop-2, showed a significant reduction in the uptake of transferrin (Tf) (**Figure 4a**,**c**) as well as GCQDs (**Figure 4b**,**d**) in both kidney and liver primary cells. (**Figure 4 and supplementary information, figure. S5a**,**c**) whereas, upon treatment with lactose, transferrin (**Figure 4a**,**c**) and GCQDs (**Figure 4b**,**d**) uptake was not affected in both kidney and liver primary cells (**Figure 4 and supplementary information, figure. S5a**,**c**).

**Figure 4.**
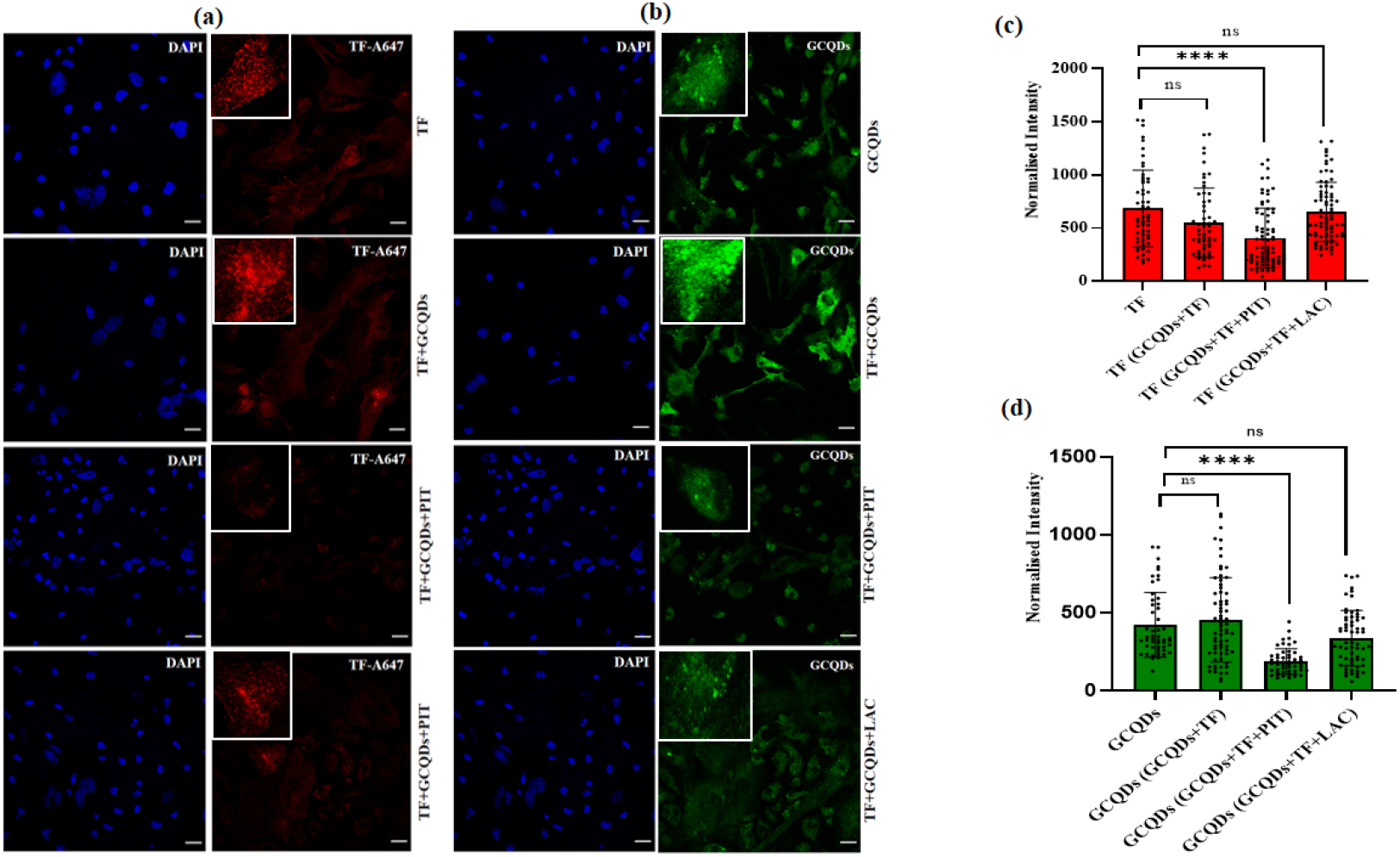
Uptake of GCQDs via clathrin mediated endocytosis in mouse kidney primary cells. **(a)** Uptake of A647 labelled transferrin (Tf) in kidney primary cells treated with pitstop-2 (20µM) and lactose (150mM) in the presence of GCQDs **(b)** Uptake of GCQDs in kidney primary cells treated with pitstop-2 (20 μM) and lactose (150 mM) in the presence of transferrin (TF). **(c)** Represents the normalised intensity and quantification analysis of Transferrin (Tf). **(d)** Represents the normalised intensity and quantification analysis of GCQDs. Scale bar is set at 20 µm of all the images. **** denotes the statistically significant p-value (p<0.0001), whereas, ns indicates statistically non-significant p-value. (one–way ordinary ANOVA). Total 50 cells and n = 2 independent experiments were performed.

Cells when treated with CIE pathway inhibitor lactose (150mM), a significant reduction in galectin-3 uptake was observed (**Figure 5a**,**c, and supplementary information, figure S6a**,**c**), while GCQDs internalization was not affected in both kidney and liver primary cells (**Figure 5b**,**d**, and **supplementary information, figure S6b**,**d**). Upon treatment with pitstop-2, galectin-3 uptake was reduced in both kidney and liver cells (**Figure 5a,c**, and **supplementary information, figure S6a**,**c**) and GCQDs uptake was only marginally reduced in kidney cells (**Figure 5b,d**), and not at all affected in liver cells (**supplementary information, figure S6b,d**).

**Figure 5.**
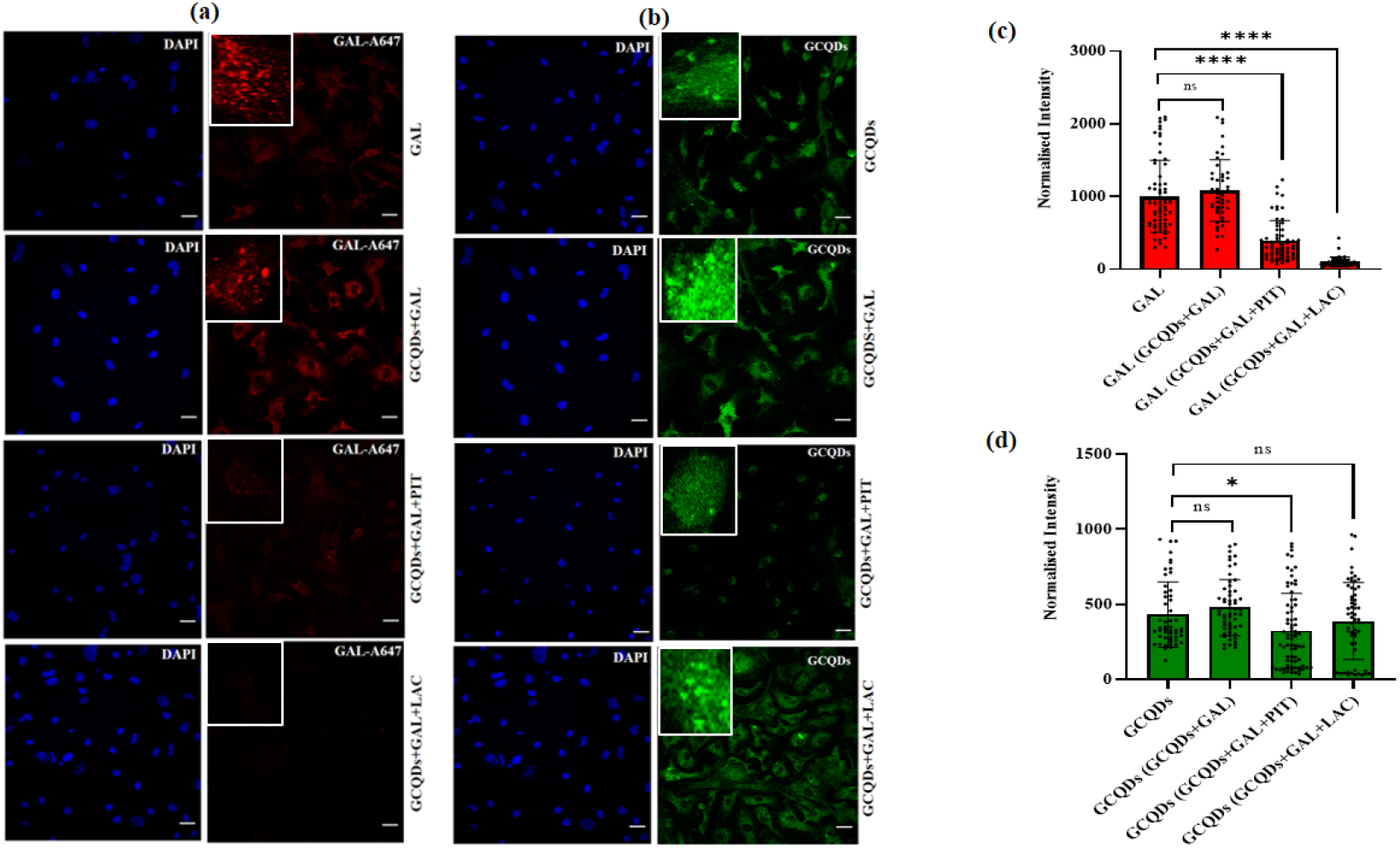
Uptake of GCQDs via Clathrin dependent Endocytosis in mouse Kidney Primary Cells. **(a)** Represents the uptake A647 labelled Galectin-3 (GAL) in kidney primary cells treated with Pitstop-2 (20µM) and Lactose (150mM) in the presence of GCQDs **(b)** Shows the uptake of GCQDs in kidney primary cells treated with Pitstop-2 (20µM) and Lactose (150mM) in the presence of Galectin-3 (GAL). **(c)** Represents the normalised intensity and quantification analysis of Galectin-3 (GAL). **(d)** Represents the normalised intensity and quantification analysis of GCQDs. Scale bar is set at 20 µm of all the images. **** denotes the statistically significant p-value (p<0.0001), * denotes the statistically significant p-value (p=0.0287) whereas, ns indicates statistically non-significant p-value. (One –way ordinary ANOVA). Total 50 cells and n=2 independent experiments were performed.

These observations establish that GCQDs uptake was significantly reduced in the presence of pitstop-2 (CME blocker) treatment and uptake was not affected with lactose (CIE blocker) treatment in both kidney and liver primary cells. Moreover, though the inhibition of CIE pathway, GCQDs is accumulated and internalized into the cells, suggesting that GCQDs is getting uptaken predominantly via clathrin mediated pathway in kidney and liver primary cells derived from mice (**Figures 4, 5 and supplementary information, figure S5**,**S6**).

### *ex vivo and in vivo* uptake of GCQDs in mice derived tissue sections and zebrafish embryos

Having established the *in vitro* cellular internalization of GCQDs via clathrin-mediated pathway, we further extended our analysis in *ex vivo* uptake of GCQDs in mice kidney and liver organ tissue sections, which represent *ex vivo* 3D models to physiologically resemble the complexity of the *in vivo* tissue environment. To understand this, we exposed GCQDs in mice derived kidney and liver tissue slices for specific time intervals, fixed and proceeded for confocal imaging to further observe the uptake of GCQDs. For the quantification, mean fluorescence intensity of each concentration of incubated sections were imaged and quantified. Surprisingly, quantification analysis revealed an uptake of GCQDs were significantly higher in kidney tissue slices as compared to control tissue slices and mean fluorescence intensity increases as the concentration increased (**Figure 6a**,**b**). Whereas, liver slices showed a significant increase in total fluorescence intensity only in 200µg/ml of GCQDs concentration (**supplementary information, figure S3**). These findings further suggest that GCQDs uptake and cellular internalization is tissue specific, preferably more in kidney tissue as compared to liver tissue.

**Figure 6.**
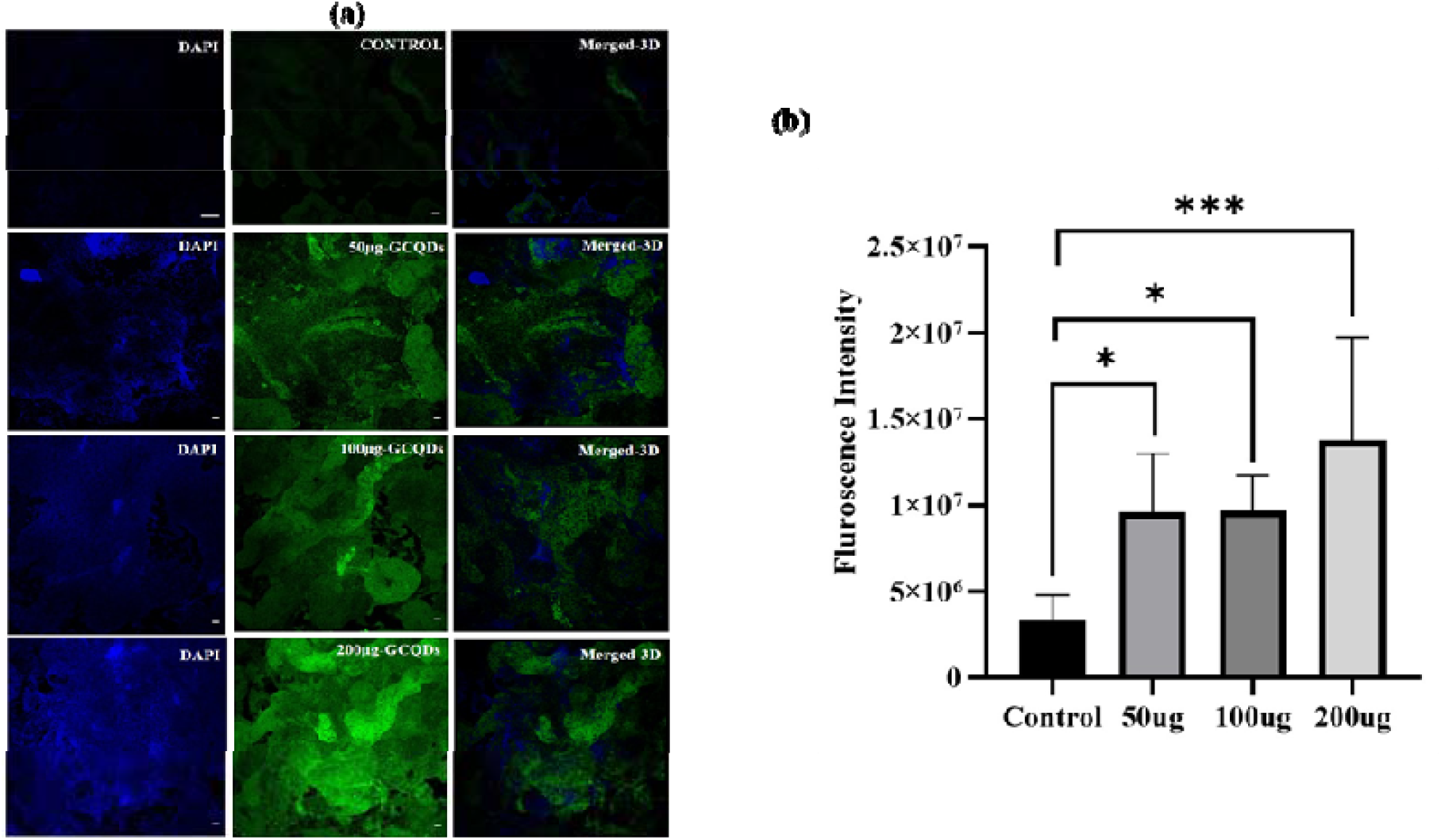
Uptake of GCQDs in kidney tissue sections. **(a)** Uptake of different concentrations of GCQDs vs. non-treated GCQDs in kidney tissue sections **(b)** Fluorescence intensity and quantification analysis of uptake of different concentrations of GCQDs in kidney tissue slices. Scale bar is set 20 µm. *** denotes the statistically significant p-value (p = 0.0004).* indicates the Scale statistically significant p-value (p = 0.0326). n = 5 tissue slices per concentration were analysed.

To explore the in vivo imaging potential of these GCQDs, the uptake of GCQDs on zebrafish larva (72 hpf) was studied by exposing the larva to GCQDs at different concentrations (100 and 200 µg/mL) for 6 h and 250 µg/mL for 12 h. GCQDs uptake was significantly higher in both 100 µg/mL & 200 µg/mL concentration as compared to control post 6 hours of incubation (**Figure 7**). Uptake of GCQDs increased with higher concentrations as the bioavailability of these nano formulations was increased to larvae. At 6h post treatments, the fluorescent intensity of GCQDs was significantly higher in the yolk sac, tail region and muscles of larva at 100 and 200 µg/mL. Despite the decreased fluorescence intensity at 250 µg/mL post 12 h treatments in larva compared to other treatments, the uptake in the yolk sac and the tail region was significantly higher than in control (**Figure 7d**). This uptake analysis highlights the bioaccumulation potential of GCQDs in developing zebrafish larva and these findings further confirm the bioimaging potential of GCQDs in zebrafish larva.

**Figure 7.**
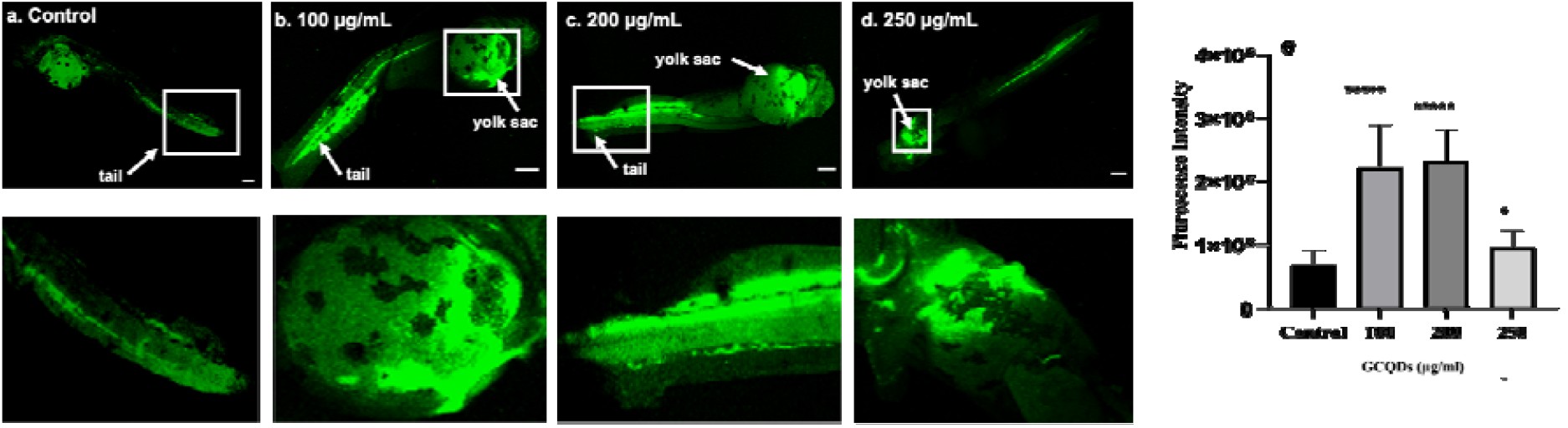
Uptake of GCQDs in zebrafish larva. **A.** Uptake of different concentrations of GCQDs in 72 hpf zebrafish larva post 6 h and 12 h treatments. (**a-c**) 6 h treatment. (**d**) 12 h treatment. **(e)** Fluorescence intensity and quantification analysis of uptake of different concentrations of GCQDs in 72hpf zebrafish larva. Scale bar is set 200µM. **** denotes the statistically significant p-value. (p < 0.0001). *denotes the statistically significant p-value (p < 0.05). 08 larva per condition were quantified.

## Discussion and conclusions

Visible spectrum emitting, bright and stable, non-blinking quantum dots available from common organic ingredients is always desirable for biomedical applications like sensing and theranostics. The organic or carbon-based materials offer a really competitive alternative to classically used inorganic nanomaterials, in that their biocompatibility, robust chemistry and most importantly the availability in large quantities. In this study, green fluorescent quantum dots (GCQDs) were synthesized using citric and ascorbic acid. The presence of alcoholic and carboxylic groups on their surface makes them highly water soluble. The cytotoxicity analysis of the GCQDs was revealed in RPE1 cells, where 90-95% of the cells were viable upto the concentrations of 300 ug/ml. This is really encouraging that the surface and chemistry of these GCQDs can be further modified to carry higher doses of biological entities or deliverables. The green emission of GCQDs was further utilized to investigate the endocytosis pathway in mouse primary cells, mouse tissues and zebrafish. GCQDs were endocytosed through clathrin mediated pathways in both mouse kidney and liver cells. The significant uptake of GCQDs in mouse liver and kidney tissues showed the potential of GCQDs for bioimaging tissues or even organs. The animal model study of GCQDs shows their significant uptake into zebrafish and can be used for targeted bioimaging in other biomedical applications. The future directions in this regard would be to improve their surface properties by protecting polymers as well as fine tuning their fluorescence emission properties and efforts to increase their stability and fluorescence quantum yields for their future sojourns into the biological universe.

## Materials and Methods Materials

Citric acid anhydrous (extrapure) purchased from SRL chemicals and L-Ascorbic acid (99.5%) from Loba chemicals. Silicon oil was purchased from X-chemicals, ethanol (>99.9%) was supplied by Changshu Hongsheng Fine Chemicals Co.,Ltd., 0.22 micrometer filter, and deionized water was obtained from Merck millipore. Lactose was purchased from Merck. The cell culture dishes dimethyl sulfoxide (DMSO) and rhodamine B were purchased from HiMedia. DMEM (Dulbecco’s modified Eagle’s medium), FBS (fetal bovine serum), and trypsin-EDTA (0.25%) were obtained from Gibco. All the purchased chemicals were of analytical grade without the need for further purification.

### Synthesis of GCQDs

1 gram of citric acid and 1 gram of ascorbic acid dissolved in a 2:1 ratio of water and ethanol. The obtained reaction mixture was then refluxed at 130ºC for 12 hours. After 12 hours the reaction mixture was cooled down naturally to room temperature. Formed green fluorescent quantum dots (GCQDs) were collected into a 50 ml falcon tube by removing the solvent by lyophilization or centrifugation.

### Characterization of GCQDs

The pH of obtained LNPs was adjusted from 1.85 to 7 using 10M NaOH. GCQDs at pH 7 were filtered using a 0.22 micrometre filter. The optical properties of GNPs, UV-Vis absorbance and fluorescence emission were recorded using Spectrocord-210 Plus Analytokjena (Germany) and FP-8300 Jasco spectrophotometer (Japan) respectively. The sample preparation for AFM was done on a freshly peeled mica sheet. A drop of filtered GCQDs at pH 7 was dropped on a mica sheet. The mica sheet was then kept in a desiccator for drying. Finally, the AFM imaging was done in tapping mode using the Bruker AFM instrument. The sample preparation for TEM was done by putting a drop of filtered GCQDs at pH 7 on a copper grid. For further characterization for example FTIR, and powder XRD, GCQDs at pH 7 were lyophilized and their powder was obtained. FTIR spectra were recorded from 400 cm^-1^ to 4000 cm^-1^ using spectrum 2, PerkinElmer in ATR mode. The powder xrd diffraction pattern was recorded using Bruker-D8 Discover with a speed of 0.2/min from 5 to 60 degrees. The GCQDs were redissolved in water after lyophilization for bioimaging in cells, mouse primary cells, mouse tissues and zebrafish.

### Quantum Yield Calculation

The quantum yield (Φ) of the GCQDs was calculated using rhodamine B as reference. For calculation of quantum yield, a solution of concentration of 1mg/ml of rhodamine B (RB) was made in milli Q, and a further 0.1 ml of this 1mg/ml was taken and dissolved into 9.9 ml of milli Q so that final concentration of RB was 10 ug/ml. The absorbance and fluorescence were taken by taking 2 ul of RB in the cuvette keeping the final volume to 3 ml . The next reading was taken by taking 4ul of RB and keeping the final volume to be 3 ml. Similarly, the subsequent readings were taken by increasing the concentration of RB by 2 ul keeping the final volume to 3 ml. The absorbance and fluorescence were taken by taking 5 ul of GCQDs in the cuvette keeping the final volume to 3 ml. The subsequent readings were taken by increasing the concentration of GCQDs by 5 ul keeping the final volume to 3 ml.

The absorbance for both RB and GCQDs were kept less than 0.1 at 540 nm. RB (literature Φ = 0.31) and GCQDs both were dissolved in water (refractive index = 1.33) . Their fluorescence spectra were recorded at the same excitation of 540 nm. Then by comparing the integrated photoluminescence intensities (excited at 540 nm) and the absorbance values (at 540 nm) of GCQDs with the reference RB quantum yield of the GCQDs was determined. The data was plotted (**Supplementary figure S4**) and the slopes of the sample (GCQDs) and the standards (RB) were determined. The data showed good linearity having mean square deviation 0.99 (reference) and sample (0.95).

The quantum yield was calculated using the below equation

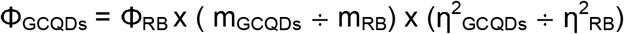

Where Φ is the quantum yield, m is slope, η is the refractive index of the solvent, RB is the reference and GCQDs is the sample. The quantum yield for GCQDs was found to be 3.3 %.

### Cytotoxicity assay of GCQDs in RPE1 cells

In order to assess the cytotoxicity of GCQDs, approximately 1×10^4^ /100µl of RPE1 (Retinal Pigment Epithelial Cell Line) cells were seeded in 96-well in media (DMEM containing 10% FBS, 1x antibiotic (Penstrep), HEPES, Sodium Bicarbonate, Sodium Pyruvate) and allowed to acclimatized for 24 h in a 5% CO_2_ at 37ºC before the experiment. Next Day, cells were washed with PBS and treated with increasing concentrations of GCQDs (50, 100, 150, 200, 250,300, 350, 400 µg/ml) for 12 hours in a serum free medium. After exposure, culture medium was discarded, 100 µl of DMEM containing MTT (5mg/ml) was added, incubated for 3-4 hours to determine the mitochondrial dehydrogenase activity of viable cells. Medium was aspirated again, and 100 µl of DMSO was added to each well to dissolve the purple formazan crystals and absorbance spectra was measured at 570 nm using Multiskan microplate reader. Experiment was done in triplicate, normalised to corresponding well containing DMSO whereas, non-treated GCQDs well was considered as control to calculate the % cell viability of each well.

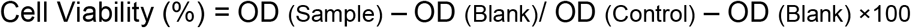

### Preparation of mouse kidney and liver primary cells

1-day old neonatal mice were rinsed off with 70% ethanol for surface sterilization and decapitated with sterile scissors. The skin was cut and the chest is open along the sternum to allow the access of the chest cavity to open the complete body to extract the liver and kidney organ tissues. Immediately, extracted organ tissues were placed and washed with sterile cold PBS (without Ca^+2^ and Mg^+2^), chopped into small pieces and digested in digestive enzyme (1x Collagenase D) at 37ºC for 20 min. Digested tissues were passed through sterile cell strainer and strained single cell suspension was centrifuged (4ºC) at 2000rpm for 10mins. Collected pellets were washed with PBS, resuspended in a prepared DMEM: F12 (1:1) medium and allowed to grow them in a flask until it reaches a proper confluence.

### Cellular Uptake of GCQDs in kidney and liver primary cells

In order to investigate by which pathway GCQDs internalised into the cell, a clathrin mediated (CME) and clathrin independent (CIE) pathways were explored in prepared mice kidney and liver primary cells. Confluent cells were trypsinized and seeded in 24-well on coverslip with density of 5×10^4^ cells, incubated with pathway specific positive marker Transferrin (5mg/ml) and Galectin-3 (5mg/ml) alone and along with increasing concentration of GCQDs (50, 100, 200 mg/ml) for 30 min in a 5% CO_2_ at 37ºC to investigate the CME and CIE pathways, respectively. Further, to probe the endocytic pathways involved in GCQDs internalization, GCQDs were incubated with CME and CIE pathway specific blocking inhibitors pitstop-2 (20µM) and lactose (150 mM) in a serum free media for 30 mins in a 5% CO_2_ at 37ºC. For the CME pathway, initially only pitstop-2 (20 µM) treatment was given to the cells for 10 min to block the ongoing endocytosis in the cells and again pitstop-2 and GCQDs combined treatments were given to the cells to assess the overall uptake of GCQDs. The untreated cells were considered as blank. Once the treatment is completed, cells were washed thrice with PBS and fixed with freshly prepared 4% paraformaldehyde (PFA) for 10 min at 37ºC. Cells were again given wash of PBS for three times and coverslips were mounted using mounting solution mowiol containing Hoechst dye for further confocal imaging.

### *Ex vivo* Cellular Uptake of GCQDs in mouse kidney and liver tissues sections

The protocol to isolate kidney and liver sections from mice are described above. From the isolated kidney and liver tissues, small sections were extracted and washed with cold PBS, immediately, tissues were snap frozen using liquid nitrogen and cut into 1 mm of small slices using sterile scalpel. Once cut, tissue slices were allowed to revive in prepared medium DMEM:F12 (1:1) for 20 min in a 5% CO_2_ at 37ºC from the shock of snap freezing. Next, the tissues were incubated in PBS for 5 min and washed twice, cut into further small slices using scalpel and incubated with GCQDs (50, 100, 200 mg/ml) for 60 min in a 5% CO_2_ at 37ºC in a serum free media (DMEM+F12). After incubation, slices were washed thrice with PBS to remove extra GCQDs and fixed with fixative solution (4% PFA) for 15 min at 37ºC. Once tissue slices were fixed, washed with PBS and mounted with Mowiol containing Hoechst for further confocal imaging.

### Animal Maintenance and Ethical Approval

1-day old Swiss albino (Mus musculus) neonatal mice were used in study. Following conditions were kept to maintain the animals. Mice pups were kept in a clean environment with 12hours light/12hours dark cycle conditions. The air was conditioned at 21±3 ºC and the relative humidity was maintained between 30-70% with 100% exhaust facility. Institutional ethical approval was obtained for all the animal experiments conducted in the study (SBR/M3/008/2022).

### Zebrafish husbandry and maintenance

The Assam wild type zebrafish were acquired from local vendors and were maintained at controlled laboratory conditions in Ahmedabad University according to the method of Kansara et al., 2019 ^20^. The male and female fishes were retained in 20 L tanks with internal conditions maintained as per ZFIN in artificially prepared fresh water. The quality of the water in the aquarium were checked regularly for the pH (6.8-7.4), conductivity (250-350 mg/L), TDS (220-320 mg/L), salinity (210-310 mg/L) and dissolved oxygen (>6 mg/L), using multi-parameter instrument (Model PCD 650, Eutech, India). The strips purchased from Macherey-Nagel Inc. (USA) were used to check the contents of ammonia (QUANTOFIX®; Reference number 90714) and nitrate/nitrite (QUANTOFIX®; Reference number 91313). The photoperiod was maintained to 14 h light/10 h dark cycle at 26-28^0^C. The zebrafish were supplemented with brine shrimp daily. Plastic traps for embryo collection were set up in the evening with the ratio of 3 females and 2 males. For experiments, embryos were collected and incubated in E3 medium (5 mmol/L NaCl, 0.17 mmol/L KCl, 0.33 mmol/L CaCl_2_, and 0.33 mmol/L MgSO_4_, dissolve in dH_2_O, adjust pH at 7.2 and autoclave E3 media; store it at room temperature) at 28^0^C.

### *In vivo* uptake of GCQDs in Zebrafish Model

*In vivo* uptake assays were performed according to organization for economic cooperation and development (OECD) guidelines. At 72 hpf (hours post fertilized), dead larva was removed and the remaining larva placed in six-well plates (Corning, NY, USA) with 15 larvae in each well. Two groups of larvae were treated with green fluorescent CQDs at concentrations of 100 and 200 µg/mL and incubated for 6 hours. One well designated as control in each group without nanoparticles. Post treatment, the medium was replaced with fresh E3 media and larva were washed to remove the excess GCQDs and fixed with fixative solution (4% PFA) for 2 minutes. Post fixation, the larva was mounted with mounting solution Mowiol and allowed to dry for further confocal imaging analysis.

### Confocal Imaging and Processing

The confocal imaging of fixed cells (63x oil immersion) and fixed tissues/embryos (10x) was performed using Leica TCS SP8 confocal laser scanning microscope (CLSM, Leica Microsystem, Germany). Different fluorophores were excited with different lasers i.e., for Hoechst (405nm), GCQDs (488 nm), Tf (633 nm), Gal(633 nm). The pinhole was kept 1 airy unit during imaging. Image quantification analysis was performed using Fiji ImageJ software. For the quantification analysis, whole cell intensity was measured at maximum intensity projection, background was subtracted and measured fluorescence intensity was normalised against unlabelled cells. A total of 40-50 cells were quantified from collected z-stacks for each experimental condition.

### Statistical Analysis

Statistical analysis was performed using Graph Pad Prism software (version 8.0.2). All the data were expressed as means ± standard deviation (SD) or means ± standard error from two independent experiments. p values were calculated using one-way ANOVA and two -tailed unpaired student t-tests with 95% confidence interval.

## Acknowledgements

We sincerely thank all the members of DB group for critically reading the manuscript and their valuable feedback. Jayishnu Roy for designing the schematic figure. PY and CM thank IITGN-MHRD, GoI PhD fellowship, PY acknowledges Director’s fellowship from IITGN for additional fellowship. KK thanks SERB, GoI for National Postdoctoral Fellowship. KS acknowledges GSBTM and IITGN for postdoctoral fellowship. DB thanks SERB, GoI for Ramanujan Fellowship, IITGN, for the start-up grant, and DBT-EMR, Gujcost-DST, GSBTM, BRNS-BARC and HEFA-GoI for research grants. Imaging facilities of CIF at IIT Gandhinagar are acknowledged. Authors declare no conflict of interest.

## Author contributions

DB and PY conceived the idea and planned the experiments. PY synthesized and characterized GCQDs. CM and SD did the cellular uptake and MTT assays in RPE1 cells. KS executed the cellular uptake and mechanisms of uptake in mice primary cells and tissues. KK executed the in vivo uptake experiments in zebrafish. AK and RR provided the zebrafish and mice facility respectively. KS and PY did the confocal imaging, KS analysed the cellular imaging data. All the authors discussed the data, helped in writing the manuscript and critically reading the manuscript.

