## Supplementary information for "Green emitting carbon quantum dots (GCQDs) to probe endocytic pathways in cells; for tissue and *in vivo* bioimaging"

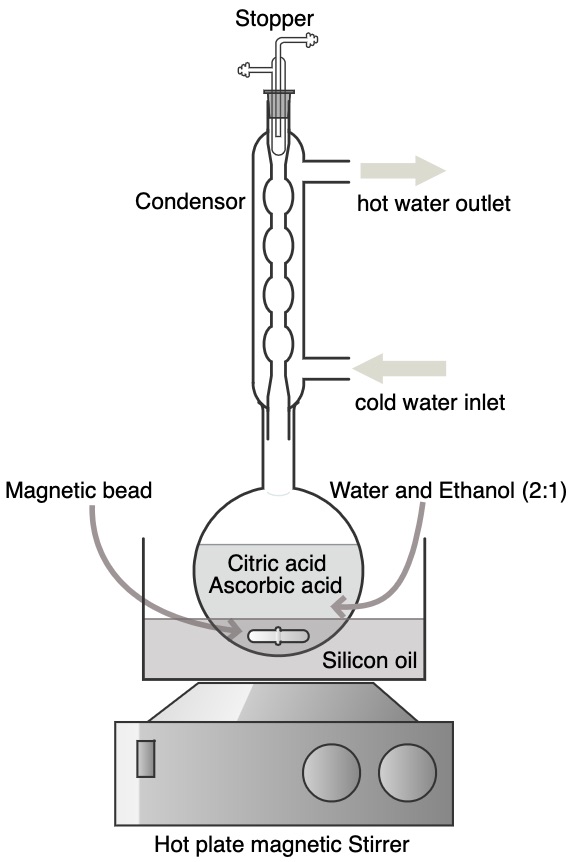

**Figure S1:** Citric acid and ascorbic acid were refluxed in solvent containing water and ethanol into a 2:1 ratio. The reaction was carried out for 12 hours at 130ºC. After 12 hours the reaction cooled down naturally and finally the harvested solution contained GCQDs.

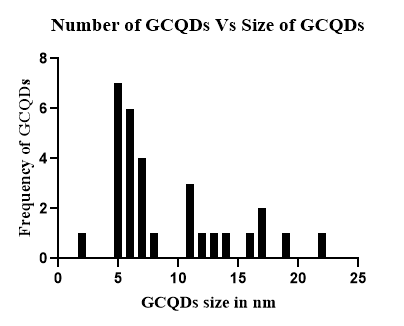

**Figure S2:** Bar graph analysis of AFM, representing number of GCQDs versus size of GCQDs. 60% (out of 31) of the GCQDs are in the range of 5-8 nm.

| 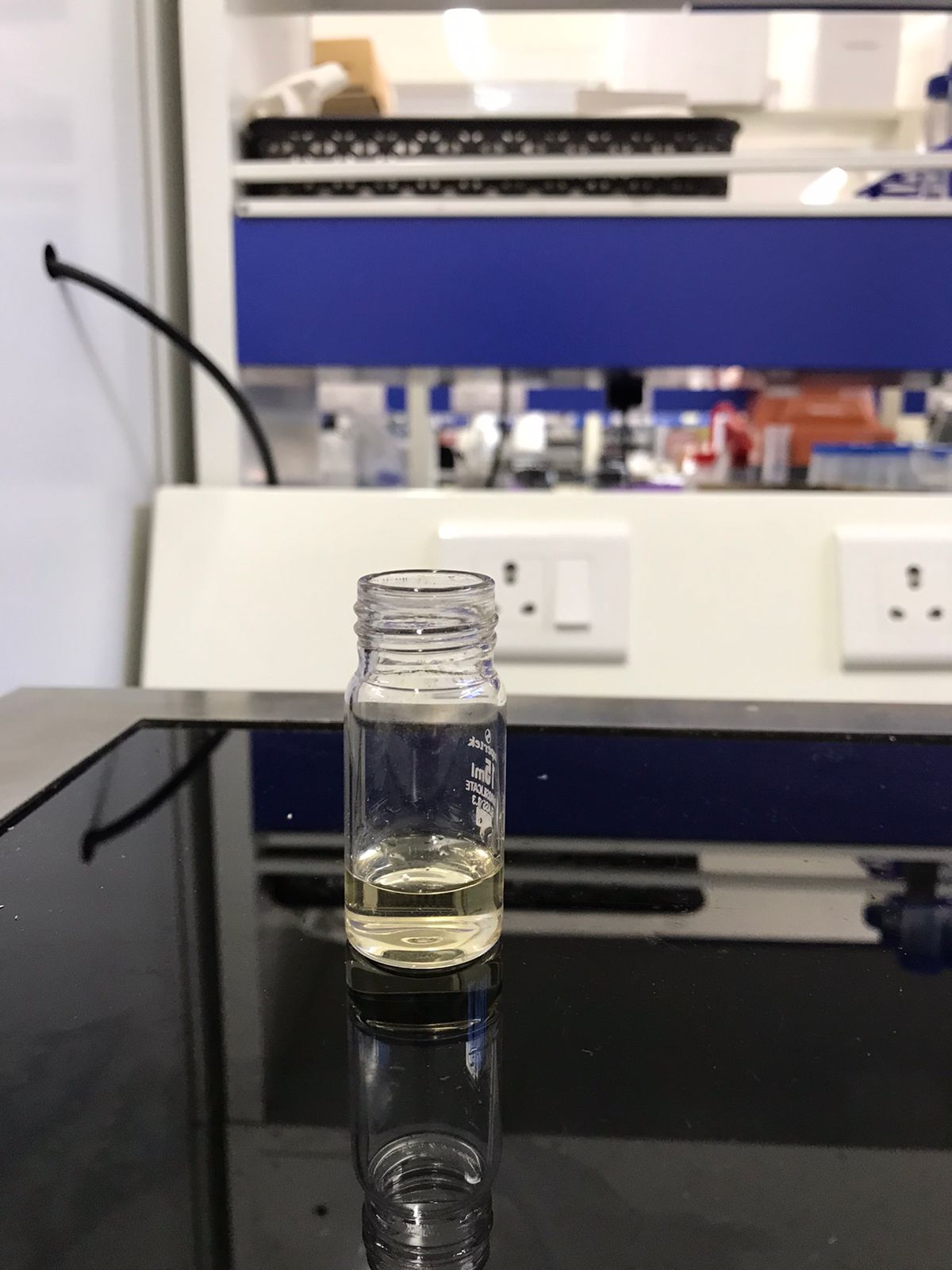 | 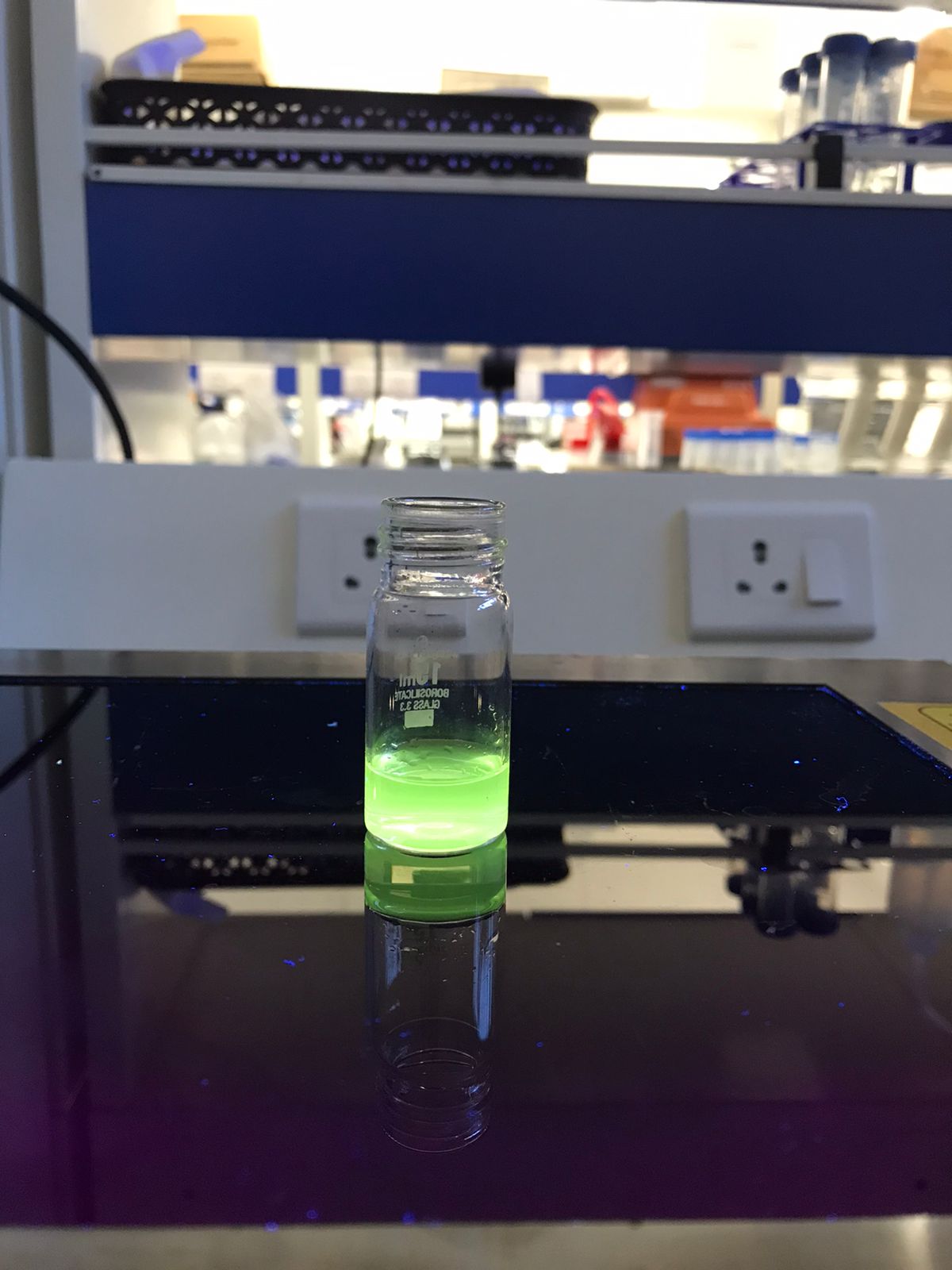 |
| --- | --- |
| **GCQDs in room light** | **GCQDs in UV light** |

**Figure S3:** The fluorescence of GCQDs in room light (on left) and under UV light (on right).

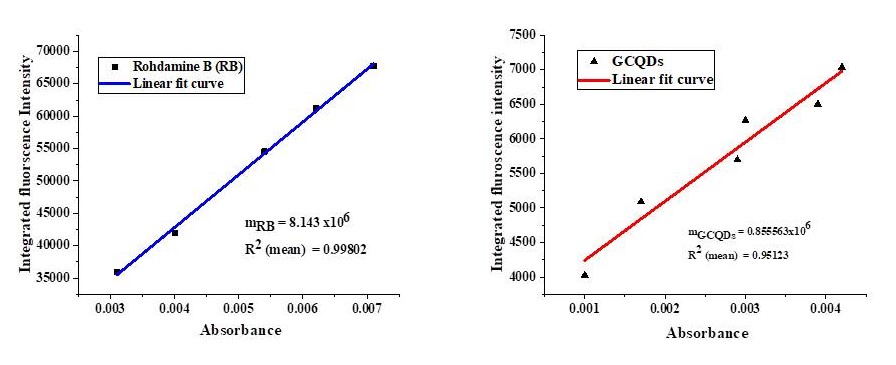

**Figure S4:** Plot of integrated fluorescence intensity of Rhodamine B as a reference (on left) and GCQDs as sample (on right) versus the absorbance. m_RB_ is the slope of reference and m_GCQDs_ is the slope of the sample. The mean square value of the best fit curve of reference and sample is 0.99 and 0.95 respectively.

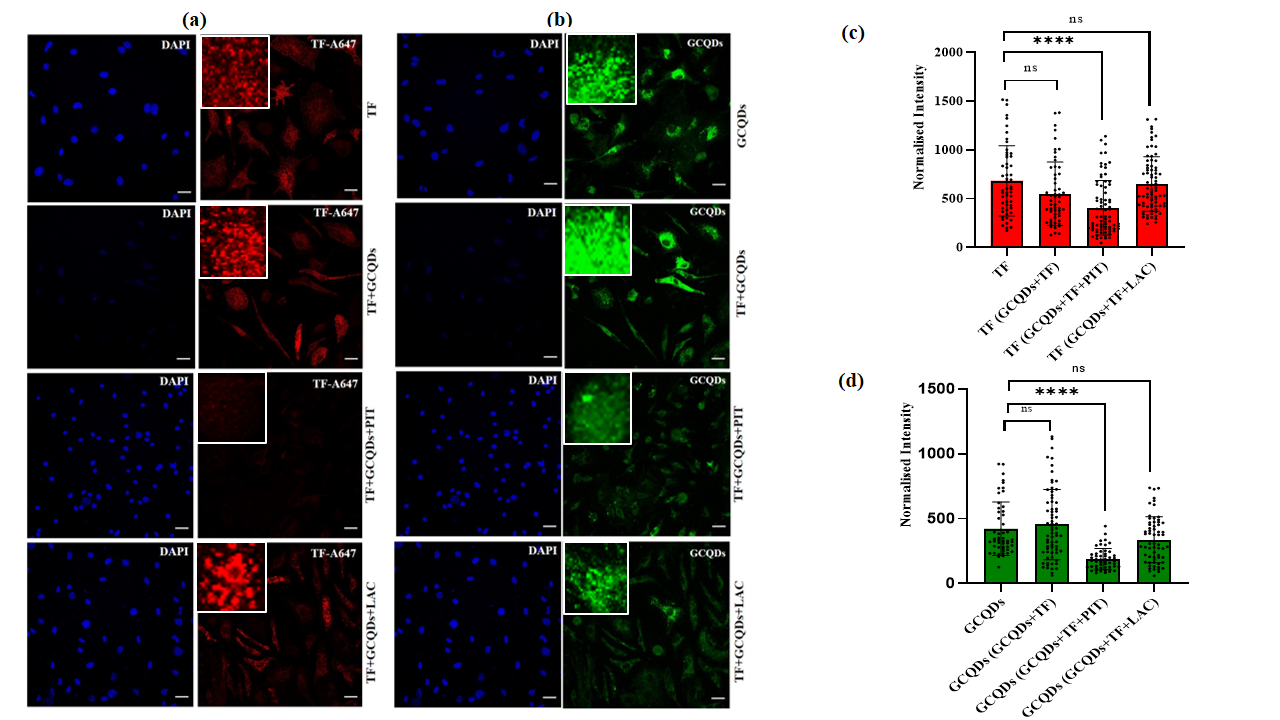

**Figure S5: Uptake of GCQDs via Clathrin Mediated Endocytosis in Liver Primary Cells.** (A) Represents the uptake of A647 labeled Transferrin (TF) in Liver primary cells treated with Pitstop-2 (20µM) and Lactose (150mM) in the presence of GCQDs (B) Shows the uptake of GCQDs in kidney primary cells treated with Pitstop-2 (20µM) and Lactose (150mM) in the presence of Transferrin (TF). (C) Represents the normalized intensity and quantification analysis of Transferrin (TF). (D) Represents the normalized intensity and quantification analysis of GCQDs. Scale bar is set at 20µM of all the images. **** denotes the statistically significant p-value (p<0.0001), whereas, ns indicates statistically non-significant p-value. (One –way ordinary ANOVA).Total 40-50 cells and (n=2) independent experiments were performed for each experimental condition.

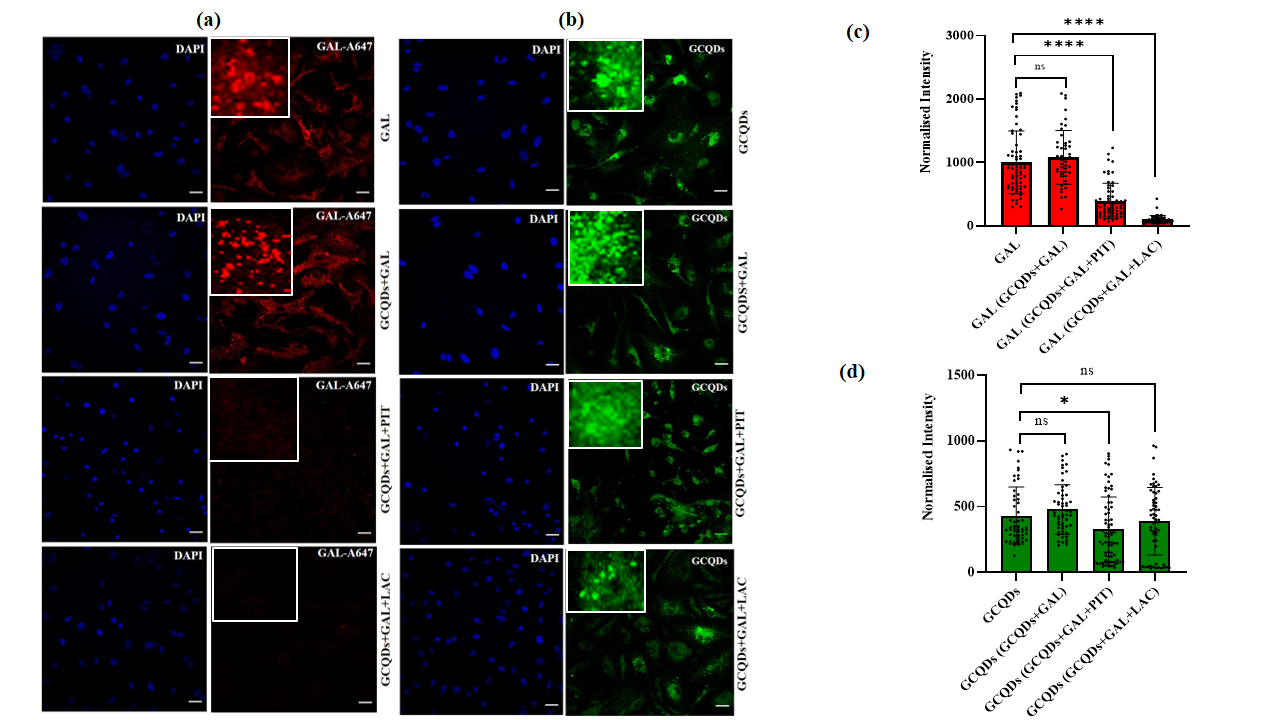

**Figure S6: Uptake of GCQDs via Clathrin dependent Endocytosis in Liver Primary Cells.** (A) Represents the uptake A647 labeled Galectin-3 (GAL) in Liver primary cells treated with Pitstop-2 (20 µM) and Lactose (150 mM) in the presence of GCQDs (B) Shows the uptake of GCQDs in kidney primary cells treated with Pitstop-2 (20µM) and Lactose (150mM) in the presence of Galectin-3 (Gal3). (C) Represents the normalized intensity and quantification analysis of Galectin-3 (Gal3). (D) Represents the normalized intensity and quantification analysis of GCQDs. Scale bar is set at 20µM of all the images. **** denotes the statistically significant p-value (p<0.0001), whereas, ns indicates statistically non-significant p-value. (One –way ordinary ANOVA). Total 40-50 cells and (n=2) independent experiments were performed for each experimental condition.

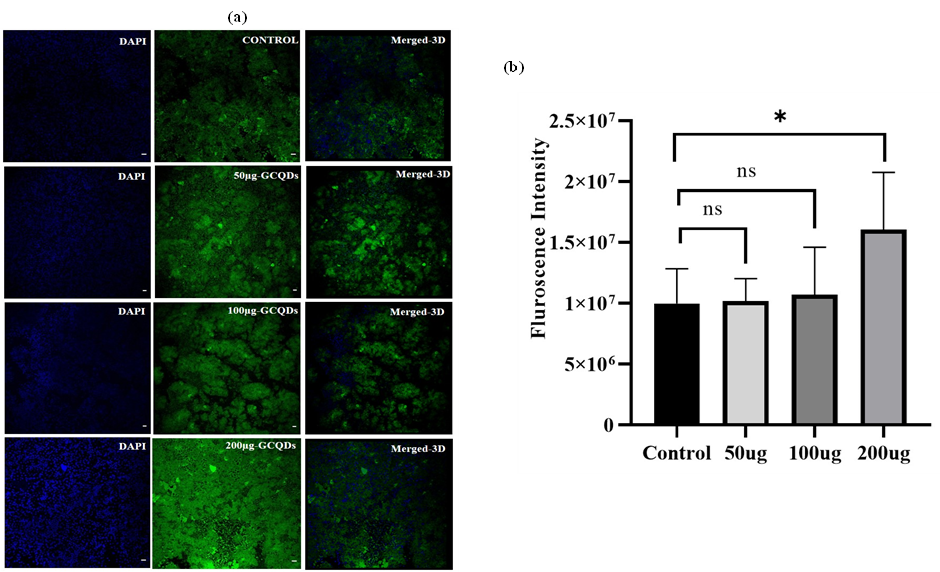

**Figure S7: Uptake of GCQDs in liver tissue slices.** (A) Shows the uptake of different concentrations of GCQDs vs. non-treated GCQDs in Kidney tissue slices (B) Fluorescence intensity and quantification analysis of uptake of different concentrations of GCQDs in Liver tissue slices. Scale bar is set 20 µM. * denotes the statistically significant p-value (p=0.0445). (n=5) tissue slices per concentration were analyzed.
